# Physiological and anatomical leaf acclimation of understory trees subjected to a through-fall precipitation exclusion in a temperate rain forest in southern South America

**DOI:** 10.64898/2026.09.01.748150

**Authors:** Ben Castro, María. F. Pérez

## Abstract

Water input is a key component of the ecosystems. Water defines the functionality, composition, and structure of biomes; therefore, any change in its availability would have an impact on ecosystem features. Moreover, a change in the water balance of an ecosystem affects its persistence as well as biochemical cycles, such as the carbon and nitrogen.

Forests are ecosystems structured using high amounts of water. Thus, trees, the oldest living plants, are the prime species in these ecosystems and are the main managers of this abiotic element. Trees uptake water from the soil, store it in their biomass, and exchange it with the environment through leaf stomata. They also intercepted rainfall and fog with their canopies. All of this water is also transmitted to the entire biological diversity that inhabits these ecosystems. Any change in water input affects the web described above. The ability of trees to modify their anatomy or processes, that is, to acclimate to novel climates, is of great advantage in maintaining the characteristics of ecosystems.

In this study, we took advantage of a precipitation exclusion experiment to reveal the acclimation of shade-tolerant understory trees, which will be the main component of a cold temperate rainforest in the future. We evaluated different anatomical and physiological leaf traits involved in the use of water by these species. We hypothesized that, as observed in similar experiments, species would adopt more conservative water-use strategies by adjusting their functional traits accordingly.

Contrary to our hypotheses, we found that understory tree species inhabiting this temperate ecosystem will not become more conservative when using water. In minimal, but significant differences, most of the studied species displayed traits, in the precipitation exclusion treatment, that were demonstrated to be water spender, rather than conservative. We attributed these contrasting changes to root metabolism alleviation due to the flooded soils of Chiloé inhabited by these forests.

## Introduction

Water is a key component for ecosystems. The changes in is availability are of high importance for the maintenance of functionality and persistence of plant communities (Bernacchi & VanLoocke 2015, Brodribb et al. 2020, Hammond et al. 2020). Plant processes, as photosynthesis, xylem increment and nutrient acquisition and mobility depends on water inputs (Taiz and Zeger 2002, Cramer et al. 2004, Chaves et al. 2008, Lambers et al. 2008), thus a change on this factor would have consequences on the economy and ecology of individuals, species and ecosystems (Vitousek 1994, McDowell et al. 2008, Seddon et al. 2016). Accordingly, processes as carbon sequestration, biogeochemical cycles and human services are linked to ecosystem water inputs (Austin et al. 2004, Chen et al. 2013, Battin et al. 2023), for instance as rainfall, fog, snow or deep aquifers. The changes in any of these components, already predicted and evidenced (IPCC 2021), would have unknown effects on earth biomes (Steffen et al. 2011). Forests, ecosystems where key plant species can live up to forty centuries, are the premier witnesses of these changes because of their long lives. Therefore, long-lived speciesneeds to possess skills, strategies, and endurance to deal with a possible change in rainfall, if its individuals wants to be successful beings (e.g. LaMarche Jr 1969, Urrutia□Jalabert et al. 2015). Depending on the rainfall shifts that a region will experience, and the plant species features, variations in structure, composition, and functionality are expected to occur, based on each specific forest. Currently, forests worldwide have experienced climatic shifts that have influenced their ecosystem characteristics (Allen et al. 2010, Hammond et al. 2022). Tree populations that have faced decreases in rainfall suffered reductions in xylem increment (e.g., Venegas et al. 2020), decline (Jiménez-Castillo et al. 2020), and more important dieback and mortality (Allen et al. 2015, Rodriguez-Catón et al. 2019, Powers et al. 2020). Temperate forests at high latitudes that have considerable inputs of water but possess a marked cold season, are greatly extended in the northern hemisphere, but they have significantly reduced distributions in the south of the equator (Armesto et al. 2001, Gilliam 2016). Southern temperate forests, including South America, are substantially different from their northern analogs, not just in extension, but also in composition and biogeography (Alaback 1991, Box 2002). For instance, the dominant species are not conifers but broad-leaved trees. Angiosperms that dominate the southern temperate forests are mostly evergreen species, while those in the north are deciduous (Axelrod 1966). Finally, species in the south are vastly endemic and descendants of the old Gondwana continent (Segovia and Armesto 2015). Consequently, they are unique ecosystems that will respond in a specific way to precipitation shifts in the future. At the early 2000’s, the Chilean National Environmental Commission formulated the “Study of the Chilean climate variability for the XXI century”. Among its results, the research concluded that summer precipitation of southern Chile will be reduced by nearly 40% by the end of the century (Fuenzalida et al. 2006). Further studies on climatic predictions agree about precipitation reductions in the region (e.g. Marengo et al. 2011, Marquet et al. 2011, Pliscoff et al. 2012, Gutiérrez et al. 2014, IPCC 2021). If environmental conditionsare maintained, and predictions meet, it would imply drastic changes for this already threatened southern South American forests. Understanding how these tree species cope with smaller amounts of water input and water stress are fundamental for the conservation and persistence of this unique temperate ecosystem. Studies addressing these last topics are new (e.g. Fajardo and Piper 2021) and most information has been generated in the west of the Andes. Dry events connected to ENSO have propitiated the dieback of *Nothofagus pumilio* in this zone of Southern south America, and the difference between dead and survival individuals have been studied (Rodriguez-Catón et al. 2016, 2019). Studies in the east of the Andes have been focused in the millennial conifer *Fitzroya cuppresoides* (Urrutia-Jalabert et al. 2015, 2018) and *Nothofagus* species (Peri et al. 2009, Piper 2011, Bucci et al. 2012). Among the angiosperm evergreen trees of the cold temperate forest, Figueroa et al. (2006) evidenced the physiological differences among *Eucryphia cordifolia* populations in the use of water. Lusk and Jimenez (2007) showed the differences in water movement rates among evergreen tree species, evidencing in some way, the fast-slow hidrological niche of each (Oliveira et al. 2021, Castro & Pérez. in revision). Although, field studies evaluating tree species acclimation lack on this part of the world (but see Bucci et al. 2019, Fajardo and Piper 2021). In this study, we measured and analyzed leaf anatomical and physiological traits in eight evergreen tree species in a temperate forest of southern South America that were subjected to a precipitation exclusion experiment and were compared to reference individuals. In the leaves, we examined fixed-kind and plastic-kind traits directly linked to water use, that also correlate to plant hydraulics, economy and gas exchange. We measured stomatal density, a trait related to the amount of gas exchanged by the plant, therefore relates to photosynthesis and stomatal conductance (e.g. Tanaka et al. 2013, Harrison et al. 2020). An increment, positive or negative, of the number of stomata in the leaf would be an indication of novel water use (Bertolino et al. 2019). We measure then leaf mass area, wish is a highly labile trait that easily can be modified to take advantage of new environmental conditions. Usually thiner-wider leaves are associated to spender individuals. On the contrary, thicker-narrower leaves are associated to a more water conservative stand (Wright et al. 2004, Reich 2014). We measure as well the economic traits, nitrogen and carbon content, which are economic indicators linked to photosynthesis and the capability of moving nutrients, therefore water usage. We also measure one of the carbon isotopes, C^13^. This trait is an indicator of stomata functioning; water use efficiency (WUE) and transpiration (Farqhuar et al. 2007), because pores that remain closer longer accumulates more of this molecule (conservative), compared to plants that maintained stomas open for longer periods (spenders). Its has been demonstrated as an adaptable trait to water scarcity conditions (Brendel and Epron 2022) and also to water availability acclimation (Domingues et al. 2018). Finally, we characterize pressure - volume curves traits. Osmotic potential at full cell turgor and osmotic potential at zero cell turgor are iconic traits that relate to hydraulics and tree species drought resistance (Barlett et al. 2012). At ecosystem and population levels, these traits usually become more negative as water availability diminishes. Relative water content at zero turgor and cell wall elasticity also provides insights of physiological water use and its adaptation and acclimation has been observed before (e.g. Joly and Zaerr 1987, Fan et al. 1994, Binks et al. 2016a). In the following study, we hypothesized that individuals that have been exposed to a diminished rainfall input will acclimate foliar traits from a spender behavior to a water conservative position. Thus, we tested the hypothesis at 1) ecosystem, 2) species and 3) traits level.

## Methods

### Study site

The study was conducted at the Senda Darwin Biological Station (SDBS), a private protected area of 120 hectares located in the north side of the largest island of Chiloé Archipelago (c. 42 LS, 73 LW, fig.1), Chile. The site is characterized by an annual rain of 2.200 mm. and mean annual temperature of 12.4 C° (SDBS meteorological station, 21 years record). Koppen and Geiger (1936) described the climate as a warm temperate climate without a dry season. During summer, the time of maximum temperatures, precipitation is approximately 300 mm. During autumn-winter, while more than 80% of precipitation falls, minimum temperatures are recorded (Frene et al. 2021).

**Figure 1.**
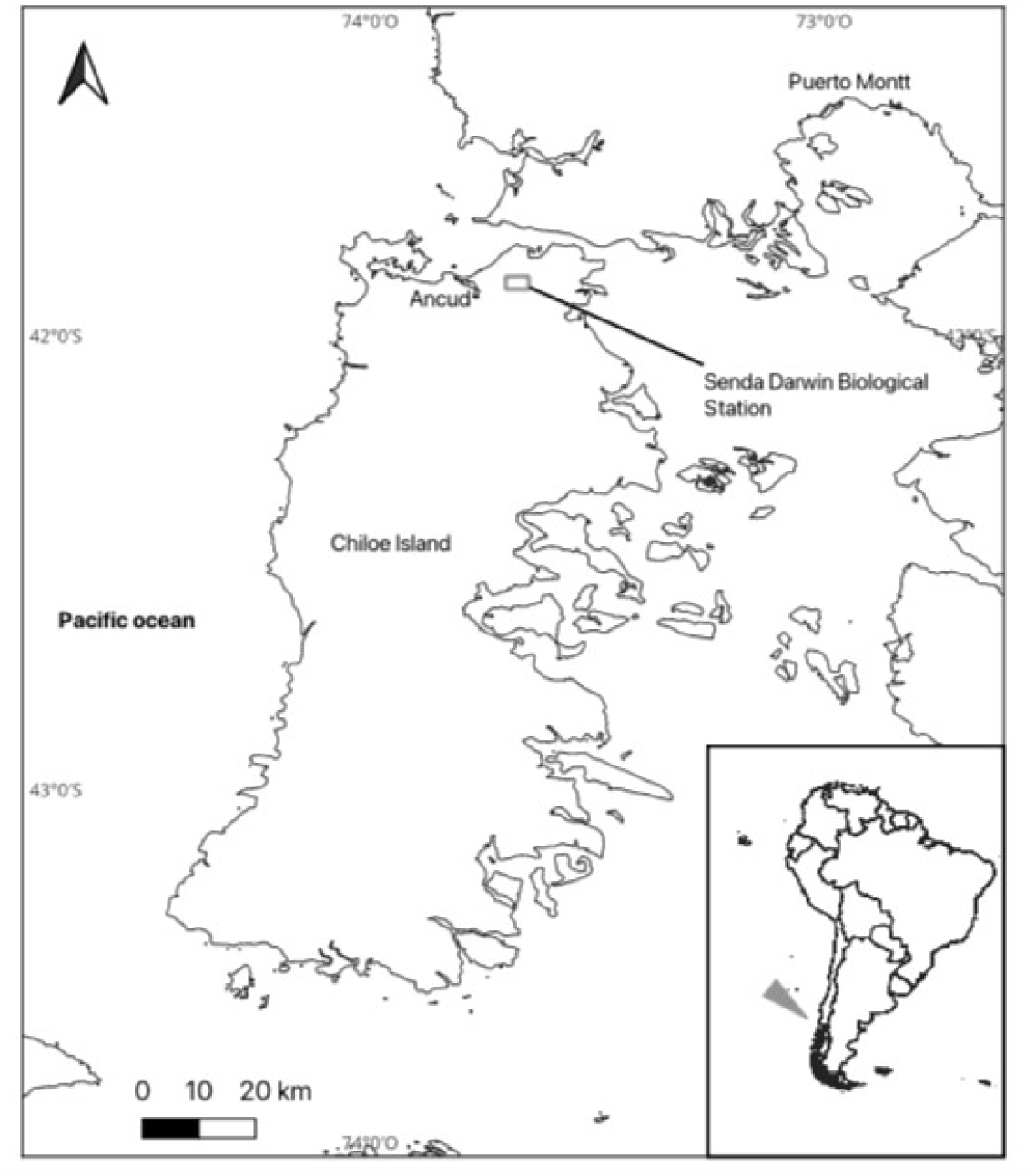
Localization of Senda Darwin Biological Station.

The geography of the northern part of the island is depicted by low hills and low elevations produced by moraine during the last glaciations (last c. 10.000 B.P.). The soils are thin and subjected to seasonal water-logging. A hardpan layer composed of iron and aluminum oxides at c. -50 cm. is responsible for the high levels of water in these soils. (Diaz et al. 2007).

Vegetation can be described as a Temperate laurophyl rainforest. The occurrence of these forest types in cool-marine areas are exclusive of the southern hemisphere (Box 2002). The ecosystem studied is a secondary forest dominated by the pioneer species, *Nothofagus nitida* and *Drimys winteri*. Individuals of both species were c. 90 years old, that form a higher canopy level at a height of 25–30 m. In the understory inhabit several species, which will become the dominant vegetation as the forest matures into old-growth—just as they probably were, around 100 years ago. These forests were completely logged for timber extraction and cattle introduction a century ago. We therefore selected the current understory species for this study. The understory is represented by shade-tolerant, evergreen species, the angiosperms trees *Amomyrtus luma, Amomyrtus meli, Tepualia stipularis, Luma apiculata, Weinmannia trichosperma, Eucryphia cordifolia, Caldcluvia paniculata* and *Gevuina avellana*. Other species as C*rinodendrum hookeranium, Lomatia hirsuta, Laureliopsis phillipiana* and *Aextoxicon punctatum* are present in SDBS, although are less frequent. This forest also hosts the gymnosperms, *Saxegothaea conspicua, Podocarpus nubigenus and Pilgerodendron uviferum*. Most of the species named above possess long-lasting lives, some of them reaching up to three hundred years (Gutiérrez et al. 2008, 2009). Additionally, *Myrceugenia parvifolia* was the most widespread shrub inhabiting this secondary forest. To the date of this study, the older individuals of the understory tree species were approximately 40 years old and had diameters from two to seven centimeters.

### Through-fall precipitation exclusion

To exclude rainfall from the forest, two sites inside the Senda Darwin station were selected as replicates. In 2011, two 20 m × 20 m plots were covered with 20 cm. × 10 m. plastic half-pipes until 35% of the total plot area was reached. Half pipes were located inside the forest under the dominant canopy at a height of approximately 3 m. All half-pipes finish in a single pipe located at the edge of the plot, which possesses a hose that transports the excluded water far from the plot (approximately 20 m. away). From 2011 to 2017, pipes were installed during the Southern Hemisphere summer (c. December–March). In 2017, exclusion infrastructure was permanently installed in both plots. Based on the rainfall partitioning study by Frene et al. (2022), we estimated that 2.847,6 mm. were excluded from each plot (fig. 2). This amount corresponds to 12% (3%; 2011-2016, 22%; 2017-2022) of the total rainfall that has fallen in this ecosystem during the aforementioned period.

**Figure 2.**
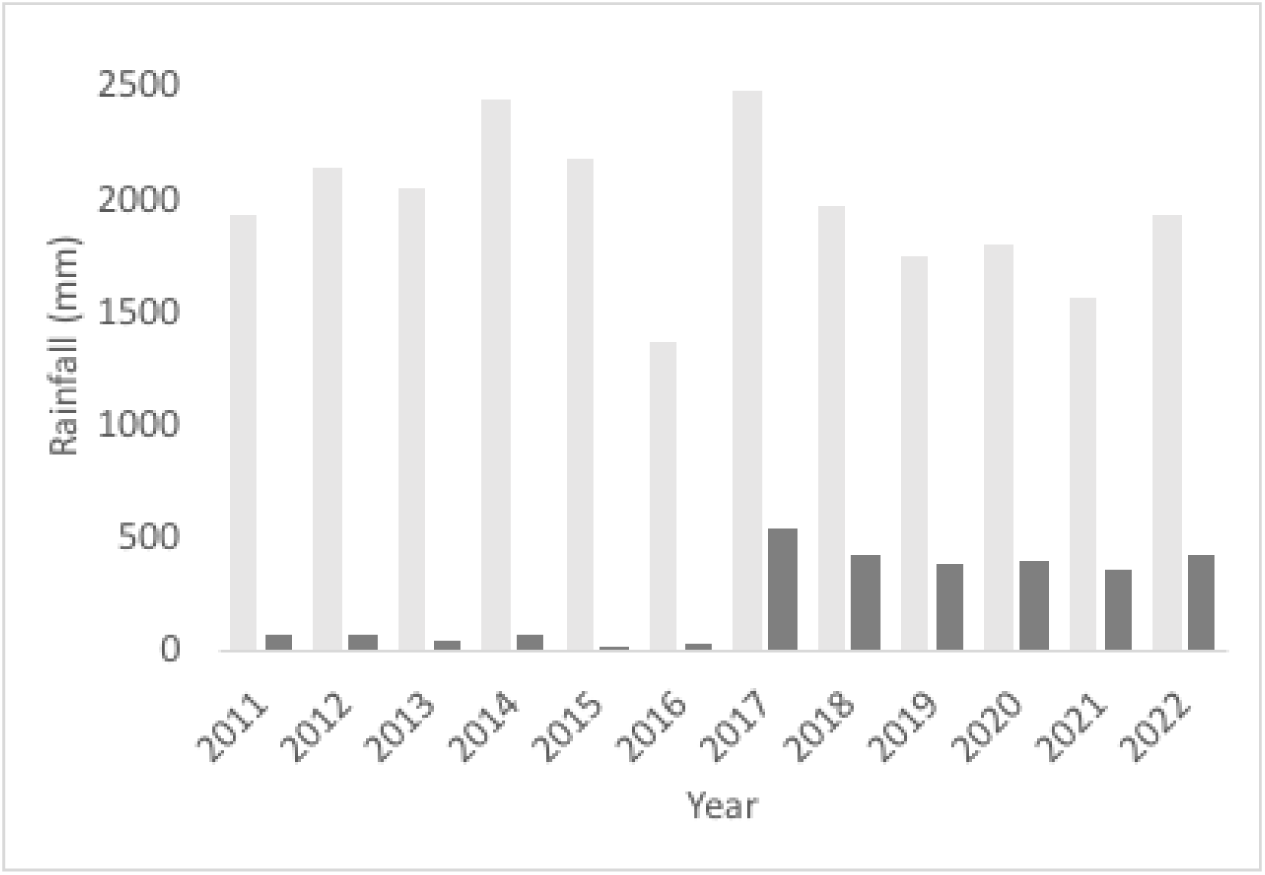
Total rainfall (light gray) and through-fall rainfall excluded (dark gray) from each plot every year since the experiment was installed in 2011.

### Species, sampling and measurement

The species included in the study are listed in Table 1. Selected species are a representative mix of the current understory forest of Chiloé. Depending on the species, we selected three to seven individuals. In a few species, there were no more individuals in the exclusion plots to investigate (*Tepualia stipularis*, 3; *Gevuina avellana*, 4). Although all the individuals of the exclusion treatment were selected inside the precipitation exclusion plots, reference individuals were selected in similar forest sites, closed to the exclusion plots. All the individuals selected have heights between 1.5 m. to 4 m. and trunk diameters from 3 to 6 cm. We sampled and analyzed 89-94 individuals, depending on the trait (42-47 in each treatment). Sampling and measurements were performed between 2021 and 2023.

**Table 1.** Species and Individuals evaluated in the study.

| Species Lineage | Species Family | Species Name | Exclusion n | Reference n |
| --- | --- | --- | --- | --- |
| Angiosperm | Myrtaceae | *Amomyrtus meli* | 7 | 7 |
| Angiosperm | Myrtaceae | *Myrceugenia parviflora* | 6 | 7 |
| Angiosperm | Myrtaceae | *Tepualia stipularis* | 3 | 5 |
| Angiosperm | Cunoniaceae | *Caldcluvia paniculata* | 7 | 7 |
| Angiosperm | Cunoniaceae | *Eucryphia cordifolia* | 7 | 7 |
| Angiosperm | Cunoniaceae | *Weinmannia trichosperma* | 4 | 6 |
| Angiosperm | Proteaceae | *Gevuina avellana* | 3 | 5 |
| Gymnosperm | Podocarpaceae | *Podocarpus nubigenus* | 5 | 5 |

### Pressure – volume curves

Pressure - volume curves (PVC) were constructed using small terminal leafy shoots of each individual assessed in the study (Table 1). Terminal shoots were collected in plastic bags, with wet paper inside to maintain moisture, and immediately brought back to the laboratory. We ensured that all leaves of the shoots were fully expanded leaves, representing at least the previous year cohort. The shoots were then cut under water and left to rehydrate in distilled water overnight.

Shoot measurements were performed using the free transpiration bench method (Barlett et al. 2012). In each fully hydrated shoot, we measured water potential and weight repeatedly, while the shoot was allowed to dehydrate during the day. Some species required more than 24 h to successfully dehydrate. Ten to fourteen measurements were performed for each individual. To measure the water potential, we used a Scholander pressure bomb (PMS 1000; PMS, OR, USA). When measurements were completed, the leaves were collected and dried for 72 h at 70 °C degrees (Venticell, Germany). Once the data were obtained, PVC was built using the protocol described by Sack et al. (www.prometheusprotocols.net). The following traits were derived from the curves: osmotic potential at full turgor, osmotic potential at turgor loss point, relative water potential at turgor loss point, and modulus of elasticity.

### Summer midday water potential

During a single day, at the end of the 2023 southern hemisphere summer (march), between 12.00 and 14.30 h. a branch with a few leaves were sampled from every individual. The leaves were immediately bagged and taken to the laboratory, where the samples were refrigerated (5°C) until measurement. The water potential of two leaves per individual was measured using a Scholander pressure bomb (PMS 1000; PMS, OR, USA).

### Leaf carbon isotopes and nutrient content

We collected 5–15 leaves per individual to measure carbon isotopes and associated nutrients. The leaves were collected and dried for 72 h at 70 °C degrees (Venticell, Germany). A sample of each leaf was then crushed, and approximately 5 mg/individual was encapsulated for measurement of carbon isotopes. An Isotope Ratio Mass Spectrometer (Thermo Delta Advantage, USA) coupled with an Elemental Flash Analyzer EA200 was used to measure the total carbon, total nitrogen, carbon to nitrogen ratio (C:N), and carbon isotopes. These measurements were obtained at the Laboratorio de Biogeoquímica e Isótopos Estables Aplicados (LABASI) from the Ecology Department of the Pontifícia Universidad Católica de Chile.

### Stomatal density

Fully expanded, preceding growing season leaves were collected, stored in plastic bags, and brought immediately to the laboratory at SDBS, and stored in a refrigerator at 5 °C. Stomata were assessed using a transparent nail paint print to obtain an abaxial impression from three leaves of 94 individuals. For *Eucryphia cordifolia*, which possesses a hairy abaxial surface, a portion of c. 0.5 cm2 was obtained from each sampled leaf and submerged in domestic chlorine for 48-72 hours. Subsequently, the cuticle was separated from the mesophyll. The epidermis was then dyed with diluted safranin 0.5% for observation (Figueroa et al. 2006)). Paint prints and *Eucryphia cordifolia* epidermis were then observed under an Olympus CX21 microscope (40X) and the stomata were counted in three different segments of the sample. The average stomata per mm2 for each individual were then obtained.

### Leaf mass area

We collected 5–15 leaves from each individual to measure leaf mass area. The leaves were then transported to the laboratory in plastic bags and stored in a refrigerator at 5 °C. To obtain the area, every leaf was scanned and then measured using J software (USA), following the traditional standard. After leaf area measurement, the samples were oven-dried for 72 hours at 70 °C (Venticell, Germany). Once the leaves were completely dried, their mass was measured using a precision scale (0.001 g, TX323L, Shimadzu, Japan). Leaf mass per area was obtained in grams per square centimeter and transformed to grams per square meter for clarity.

### Statistical analysis

We started analyzing trait-treatment-species interactions by performing a Principal Component Analysis (correlation matrix). Then, to assess the differences we performed a Two-way ANOVA using the scores of the principal components that better explain the model. Tukey test was used as post-hoc analysis. We executed this method for raw data (as measured) and mean difference data, transforming data as follow:

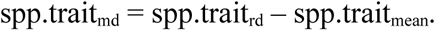

Where spp.trait_md_ is the mean difference data of a trait of each species, spp.trait_rd_ is the measured data of a trait of each species, and spp.trait_mean_ is the mean of each trait of each species.

We then used Two-way ANOVA to asses treatment-species interaction in single traits. We finally did comparisons by treatment using t-test. We used the software Past 4 (Germany) and Rstat (Rteam, USA) to perform analysis and make graphs.

## Results

### Multivariate analysis, trait-species-treatments interaction

Principal component analysis of raw data separated species and families through the two axes, but did not separate clearly the whole treatments (Fig. 3). Principal component 1 and 2 explained 29.6% and 24% of the variance of the model. Scores from the first component showed significant differences in the two-way ANOVA for the treatment, species and their interaction, although the second component presented differences just for the species (Table 2.). Component 2 axis separates Myrtaceca from the group formed by Cunoniaceae family with *Gevuina* and *Podocarpus.* This group exhibited higher values for P-V curve traits, while the Myrtaceae showed higher values for stomatal and carbon-related traits. Principal component analysis of trait mean differences provided clearer separation of treatments compared to raw data (Figure 4), with Components 1 and 4 explaining the group differences most effectively. In this case, ANOVA results were significant for treatments. The reference group displayed higher values for LMA, δ¹³C, and stomata, whereas the exclusion group was characterized by higher midday water potential values.

**Figure 3.**
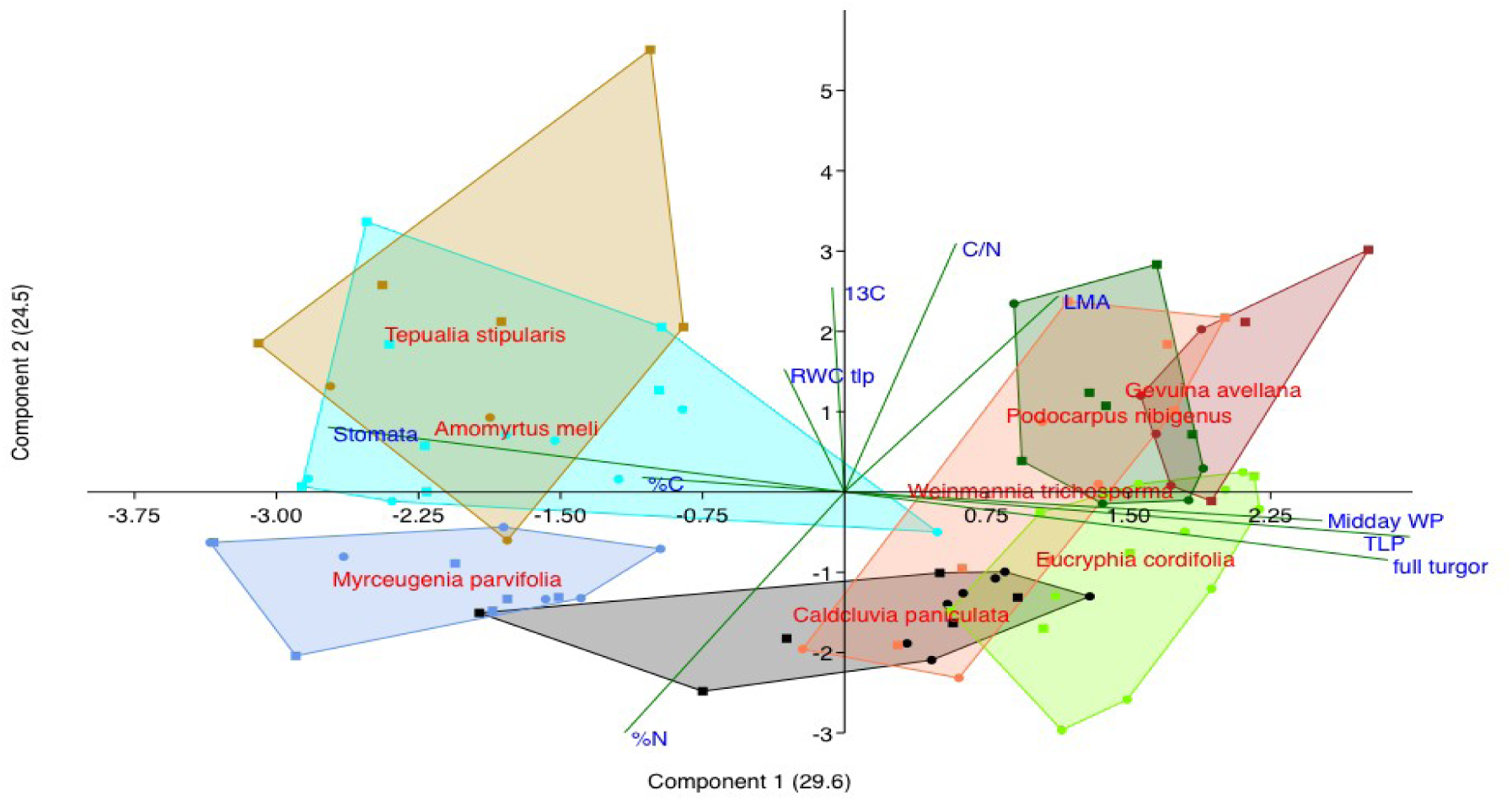
PCA of the studies individuals, using raw trait data. Polygons represent the variability of each species due to the traits assessed.

**Figure 4.**
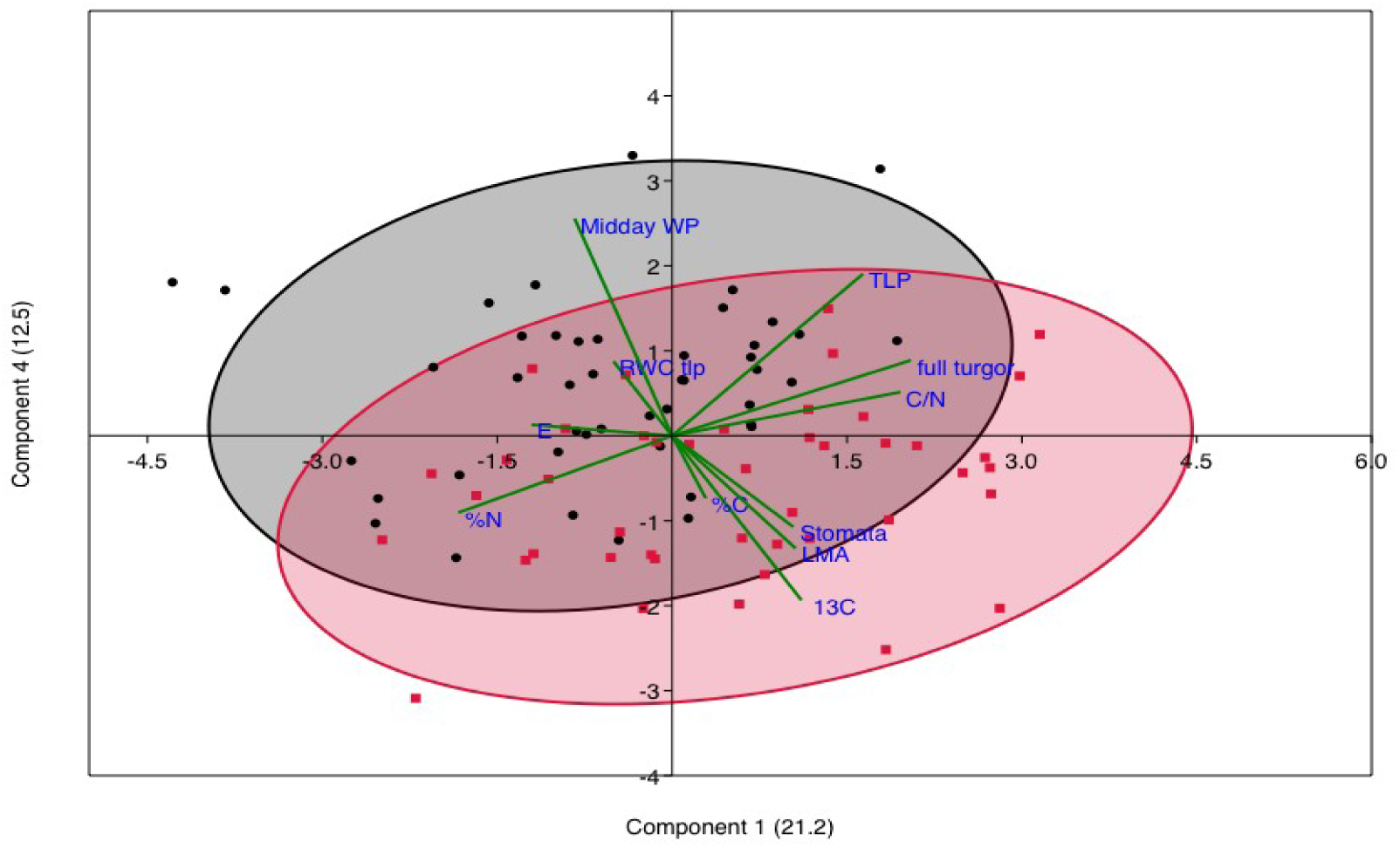
PCA using mean differences of traits. Black ellipse group is the exclusion treatment and red ellipse group is the reference.

**Table 2.** Two way ANOVA parameters of the principal components one and two.

| PC1 | Sum of sqrs | df | Mean square | F | p |
| --- | --- | --- | --- | --- | --- |
| Species: | 229.77 | 7 | 32.82 | 46.06 | 2.75E-24 |
| Treatment: | 3.07 | 1 | 3.07 | 4.308 | 0.04145 |
| Interaction: | 11.20 | 7 | 1.60 | 2.244 | 0.04002 |
| Within: | 52.03 | 73 | 0.71 |  |  |
| Total: | 292.15 | 88 |  |  |  |

| PC2 | Sum of sqrs | df | Mean square | F | p |
| --- | --- | --- | --- | --- | --- |
| Species: | 114.859 | 7 | 16.4085 | 12.72 | 1.46E-10 |
| Treatment: | 4.67997 | 1 | 4.67997 | 3.629 | 0.06071 |
| Interaction: | 14.5767 | 7 | 2.08238 | 1.615 | 0.1448 |
| Within: | 94.1336 | 73 | 1.2895 |  |  |
| Total: | 226.276 | 88 |  |  |  |

### Species treatment interactions

Most of the two-way ANOVA resulted significant for species and interactions, which it was expected from the differences in the water use strategies of these species (fig. 3). Treatment differences resulted significant for RWCtlp, midday water potential, ^δ^13C, nitrogen, C:N, LMA and stomata (fig. 5).

**Figure 5.**
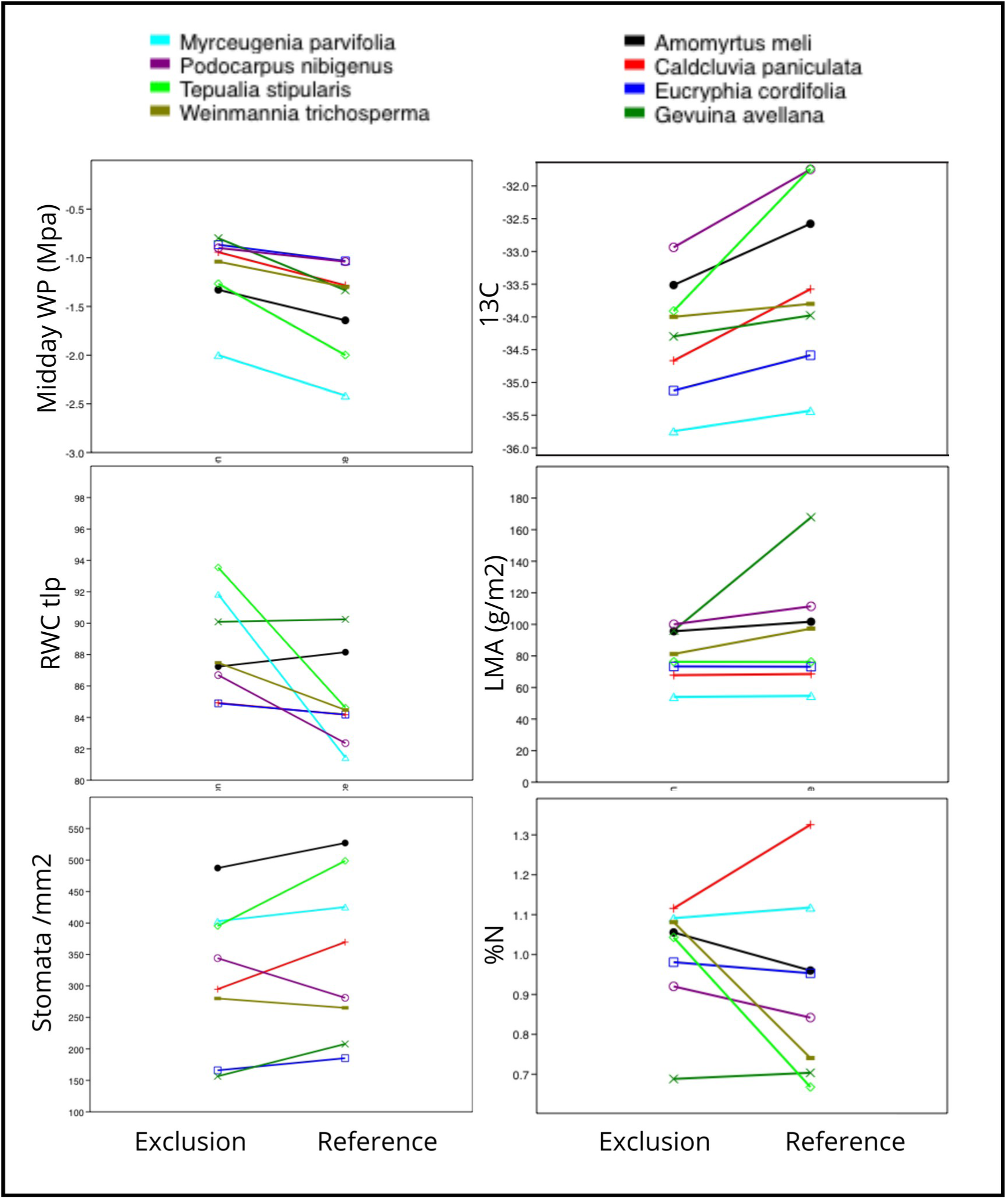
Two way Anova of the statistical significant traits-treatments by species.

### Single relationships

#### Leaf mass area relation

A greater ratio of average LMA was found in the reference ecosystem individuals compared to excluded, with the exception of *Eucryphia cordifolia*. Leaf mass area did not show significant differences when single species in both treatments were compared (Supplementary data). The average LMA was 16% higher in the reference ecosystem than in the exclusion treatment, which was more than 13 g of tissue per square metre of leaves. However, this difference was mainly driven by a few species. The Myrtaceae family did not show statistically significant differences when the Cunoniaceae family showed weak significant differences.

#### Carbon isotopes

The average ^δ^13C was consistently higher in the individuals inhabiting the reference ecosystem than in the excluded individuals (Table 3). Significant differences between the treatments were found among species and family groups. ^δ^13C was significant in *Caldcluvia paniculata, Tepualia stipularis*, and *Podocarpus nubigenus*. Significant differences for the t-test were also found for the Cunoniaceae family. Fewer strength differences were observed for the Myrtacea family.

**Table 3.**
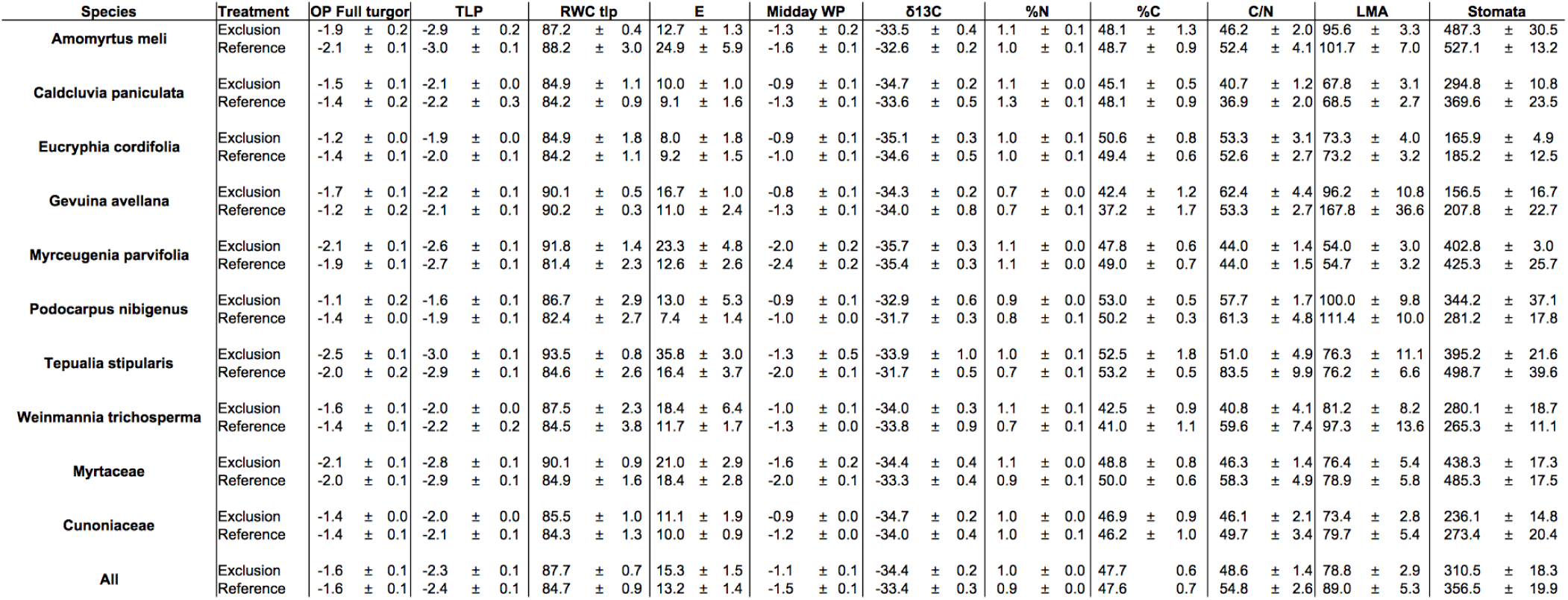
Traits evaluated of the eight species in the treatments. Average values ± standard error.

#### Nitrogen

Average nitrogen varied among species between treatments, although the content in the leaves was greater, at the ecosystem level, in the precipitation exclusion treatment (Table 2). At the species level, the difference in nitrogen content was significant for *Caldcluvia paniculata, Tepualia stipularis*, and *Weinmannia trichoesperma*. The t-test was also significant for the Myrtaceae family.

#### Carbon

Carbon resulted highly similar between treatments. Nonetheless, three species resulted with significant differences, *Caldcluvia paniculata, Podocarpus nubigenus* and *Gevuina avellana*. The Myrtaceae family showed tenuous statistical differences (Table 3).

#### Carbon to nitrogen ratio

At the ecosystem level, the C:N ratio was higher in the reference ecosystem (Supplementary data). Alongside the exception of *Tepualia stipularis*, none of the other species resulted in significant differences between treatments.

#### Stomatal density

Stomatal densities exhibited no statistical differences among the individuals of the species assessed (Appendix1). Nonetheless, the total average stomata per treatment were smaller in the excluded individuals than in the reference. A similar trend was observed for five species and both families, Myrtaceae and Cunoniaceae.

#### Pressure – volume curves parameters

Few significant differences were found for the traits of the PVC (Supplementary data); for instance, *Eucryphia cordifolia* only presented differences in osmotic potential at full turgor. At the turgor loss point, only *Podocarpus nubigenus* showed statistical differences among the treatments. The relative water content at the turgor loss point was significantly higher for one species, *Myrceugenia parvifolia*, and its family, Myrtaceae. Two of the Myrtaceae species presented differences in cell wall elasticity, although in different directions.

#### Summer midday water potential

The water potential measured in the control treatment was lower (more negative) than that in the exclusion treatment. When the average water potential of the individuals in the through-fall exclusion was -1.1 Mpa, the individuals that acts as control reached a midday water potential of -1.5 Mpa. At the family level, Myrtaceae species differed between treatments; however, Cunoniaceae species differed only slightly (Figure 6). At the species level, most species differed weakly (Supplementary data), with the exception of *Podocarpus nubigenus*, which did not show significant differences.

**Figure 6.**
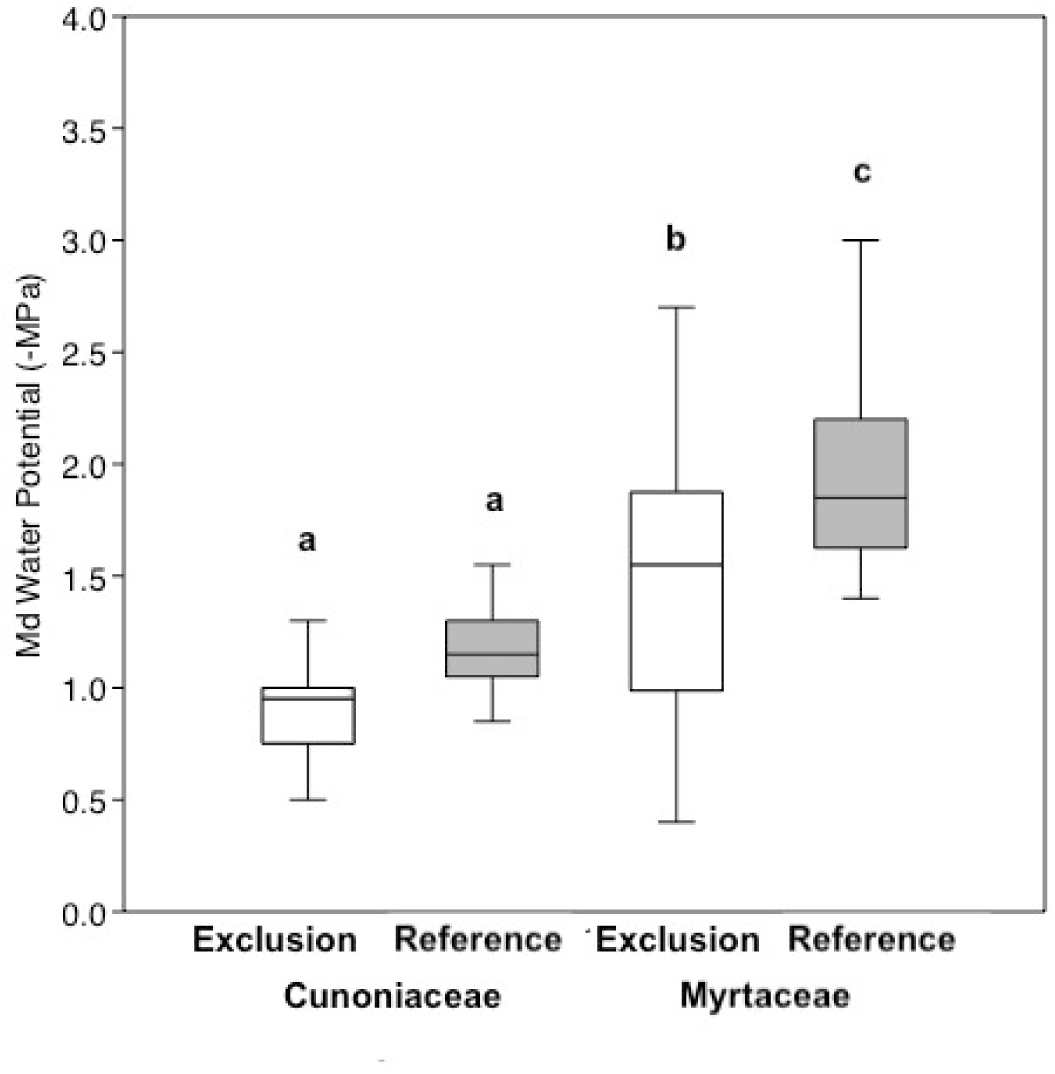
Box chart for the midday water potential at the end of summer, due main tree families and treatments. Letters represent ANOVA Tukey’s test.

## Discussion

In this study of the cold temperate rainforest of southern South America, we evaluated tree anatomical and physiological traits to understand whether a limitation of water input through precipitation exclusion would affect them, hypothesizing an acclimation to more water-conservative traits, individuals (e.g. Tng et al. 2018) and forest. We found differences at various interaction levels. Differences were found using multivariate analysis. Similarly, single leaf traits of the evaluated species showed statistical differences associated with the treatments. However, not all assessed traits differed. Moreover, results does not support our main hypothesis that traits, individuals and the forest become conservative because of reductions in water input, as has been observed in other forests submitted to rainfall exclusions (e.g. Rowland et al. 2015, Binks et al. 2016B, Bittencourt et al. 2020). In contrast, trees in the exclusion experiment appeared to become water-spender. To our knowledge, this is the first study to find trees that become spenders instead of conservatives because of an experimental reduction in precipitation (Binks et al. 2016a, Tomasella et al. 2018, Limousin et al. 2022). Contrary to our hypothesis, we found that the understory tree individuals in the exclusion experiment seemed to be slightly acclimated to employ more water than their counterparts in the reference ecosystem.

Contrary to expectations, among the traits assessed, the average ^δ^13C was always higher in the species inhabiting the reference ecosystem, that is, the ecosystem with higher precipitation exposure. Previous evidence has shown that trees subjected to lower quantities of rainfall, compared to the initial conditions, display greater levels of carbon isotope composition (for example, Picon et al. 1996, Domingues et al. 2018). Dehydrated soil with high water potentials would result in partial stomatal closure, increasing WUE and provoking a higher discrimination of the heavier carbon isotope, as seen in other studies (Leffler and Enquist 2002, Corcuera et al. 2010). Although there was a small difference in ^δ^13C between the treatments, all species in the exclusion treatment had lower levels of ^δ^13C. Significant statistical differences between treatments and tree families were found, as well as between four species. If the species in the exclusion treatment are subjected to some degree of water stress that may impair physiological functions, such as stomatal conductance or photosynthesis, leaf ^δ^13C values should be higher than those of the reference ecosystem (Farquarh et al. 1989, Dawson et al. 2002, Pflug et al. 2015). Values of ^δ^13C scarce in the southern hemisphere, with the exception of the tropical Amazon forest, and most evidence comes from North America and European forests (Shestakova et al. 2017). Furthermore, the ^δ^13C of this pool of species is lower than that of dry tropical forests, suggesting a low WUE for this set of species (Leffler and Enquist 2002). It is even lower than that of humid tropical forests (Martinelli et al. 1998), evidencing water over-abundance in this ecosystem. Accordingly, carbon isotope accumulation is lower in this ecosystem than that in the Mediterranean forest in Chile (Romay and Bown, 2021). Foliar nitrogen levels were also higher in the exclusion treatment group. Higher levels of nitrogen suggest faster carbon gain, which may be related to higher levels of stomatal conductance in the exclusion treatment, with lower inputs of precipitation (Chaves et al. 2008). Similar to ^δ^13C value differences, if individuals in rainfall exclusion are submitted to a water shortage, the nitrogen levels in the leaves should be lower (Gesler et al. 2016). Interestingly, foliar carbon content was similar between treatments, and the differences were significant only in three species. This notion makes sense if carbon limitation does not exist in this ecosystem, which is possible because of the high rainfall in this forest. Because nitrogen content in the exclusion treatment was higher, the C:N ratio also resulted in a significant difference in this trait. Among the species, C:N showed a significant difference in one species, *Tepualia stipularis*. Evidence of differences in carbon isotopes and nutrient content highlights the possibility that some tree species, and probably the ecosystem, are acquiring carbon quicker (probably allocated to root or trunk increment), transpiring more, and consequently growing leaves with more nitrogen.

More proof that contradicts our initial hypothesis that trees become water-conservative can be seen in other traits such as leaf structure. At the species level, LMA did not show any statistical differences. At the ecosystem level, the reference system had a higher level of mass per unit area than the ecosystem in the exclusion treatment (Table 3). The individuals in the non-manipulated ecosystem possess thirteen more grams of tissue per square meter of foliar area compared to the individuals in the experimental manipulated ecosystem. In terms of drought effects on the LMA, if the specimens in the exclusion treatment suffer some type of water shortage, the LMA should be higher than that in the reference ecosystem (Wright et al. 2004, Salgado-Negret et al. 2015). Thus, it seems that in this case, rather than increasing the LMA in the reference, exclusion specimens have decreased it. Nevertheless, this effect was more apparent in half of the evaluated species, and the other half did not acclimate their leaves because of lower water inputs.

We found that the average stomata per square milliliter was significantly lower in the exclusion treatment group than in the control group. For instance, *Populus* species have demonstrated a positive relationship between stomatal density and stomatal conductance, and negative relationships with WUE and ^δ^13C (Cao et al. 2011). However, stomatal density differences have been linked to both an increase and a decrease in environmental water in species of this forest (Figueroa et al. 2010, Salgado-Negret et al. 2015). However, these changes appear to be attributable to an adaptation process rather than an acclamation process. For instance, *E. cordifolia* populations have more stomata in places where more suitable photosynthetic conditions are met (Figueroa et al. 2010). In contrast, individuals of *Aextoxicon punctatum*, another inhabitant of this forest, have reduced stomatal numbers in conditions where more water is available (Salgado-Negret et al. 2015). If we consider water use strategies in this ecosystem, conservative species (Myrtaceae) possess a higher stomatal density than spenders (Cunoniaceae); therefore, a decrease in stomatal density could be a response to higher stomatal conductance in this forest.

Pressure-volume curve parameters also showed some changes that could be related to trees becoming more expenders than conservatives. The first evidence was that none of the individuals showed a significant change related to a conservation strategy because of decreased water input (Table 2). Other studies have shown that the OPFT and TLP are associated with the level of water to which a species is exposed (for example, Salgado-Negret et al. 2015), but most link it to population adaptations rather than individual acclimation. However, Binks et al. (2016b) reported differences in these traits in a through-fall exclusion experiment, where individuals exposed to reduced water availability acclimated to more negative values. For instance, *Eucryphia cordifolia* was the only species that significantly modified OPFT, diminishing it in the exclusion treatment (Table 2), indicating that more water was available for use, requiring less solute in the leaf cells of these individuals. Osmotic adjustments have also been suggested for this species (Figueroa et al. 2010). A similar situation was observed in *Podocarpus nubigenus* but with TLP. The single conifer in this study reduced the volume of solutes at the TLP in the trough-fall reduction treatment, indicating an increase in theavailable water for use. *Myrceugenia parvifolia* and the Myrtaceae family had significantly higher RWCtlp values in the exclusion group than in the reference individuals. This statement also suggests that more water is available for use in the treatment. Prior evidence shows contrary results (for example, Salgado-Negret et al. 2015, Binks et al. 2016b): species subjected to water scarcity tend to have less water at the TLP because, otherwise, they will be more exposed to rapid cessation of metabolic processes if water is scarce. Elastic adjustment was also significant in two species of the Myrtaceae family, *Amomyrtus meli* and *Tepualia stipularis*. However, this change occurred in the opposite direction. While *A. meli* significantly reduced the modulus of elasticity under exclusion, *Tepualia stipularis* increased it. Similar to stomatal density, cell wall elasticity can occur in both directions if water availability changes. For example, a species can make its cell wall more elastic so that it can store more water (Salleo et al. 1997). On the other hand, cell walls can be made more rigid, so that the stocked water can be retained more robustly (for example, Saito and Terashima 2004, Hessini et al. 2009). Cell wall elasticity is a highly variable trait that changes throughout the year, and its mechanics and trends remain elusive (for example, Abrams 1990, Zhang et al. 2019). Osmotic and elastic adjustments are divergent strategies among plants (Lambers et al. 2008), and both traits are genetically fixed. Overall, the parameters obtained in this study are in agreement with the global assessment (Bartlett et al. 2012), although the TLP seems to be higher (more negative) in this study for a few species than in previous studies (Figueroa et al. 2010, Jiménez-Castillo et al. 2011).

The differences in the midday summer water potential among the treatments are noteworthy. All species had significantly lower water potential in the exclusion treatment. This last sentence suggests water stress alleviation for individuals subjected to reduced rainfall, instead of an increase, expected from such manipulation (Bittencourt et al. 2020). Other studies have shown such a decrease in operational water potential in natural populations (Figueroa et al. 2010) and in other exclusion experiments (Limousin et al. 2022). Waterlogged soils, which are highly common in Chiloé, may cause symptoms that are similar to drought stress, including wilting, which can be reflected in the measurement of the water potential (Lambers et al. 2008).

Other published studies that address the effects of precipitation exclusion on tree anatomy and physiology have shown that trees possess a capability, although limited, to acclimate to higher water stress. For instance, Binks et al. (2016a) found that leaf morphology of amazonian trees submitted to a precipitation exclusion experiment did not acclimate to be more conservative in the use of water. Nevertheless, leaf-water relations did response to the water deficit, acclimating to the new conditions (Binks et al. 2016b) Bittencourt et al. (2020) demonstrated a low capacity to adjust hydraulic traits in Amazonian trees, although they were more able to modify their hydraulic status. Recently, Petit et al. (2022) found no xylem acclimation, but more carbon was allocated to the leaves of trees in the rainfall exclusion. This evidence demonstrates that tree species can modify their traits—albeit to a limited extent—to adopt more water-conservative strategies, which contrasts with our findings. Domingues et al. (2018) found the opposite to our results in understory species. The species in the exclusion experiment had higher amounts of carbon isotopes, suggesting a stomatal constraint. In our study, the exclusion treatment resulted in a reduced amount of leaf carbon isotopes, indicating alleviation of the gas exchange behavior. Acclimation in traits such as turgor loss point (TLP) and midday water potential has also been reported in similar studies (e.g., Binks et al. 2016b), where trees exhibited more conservative water-use strategies—characterized by the regulation of water tension traits toward more negative water potentials (Limousin et al. 2022). Higher tension in the xylem has been argued to be the mechanism of tree decline and dieback in the Amazon (Bittencourt et al. 2020). In contrast, we observed that the individuals acting as controls presented more negative midday water potentials. We measured midday water potential at the end of summer, immediately after the driest and warmer period, probably representing the maximum differences between the individuals in both treatments. In addition, is important to note that most of these evidences come from tropical forests, and small evidence exists in temperate broadleaved evergreen forests. For instance, Schuldt et al. (2011) observed changes in foliar traits, but they did not observed differences in ^δ^13C. The super-humid ecosystem evaluated in that study may be more similar to our forest than to other tropical rainfall exclusion experiments, which have more marked dry seasons (for example, Neptad et al. 2007, da Costa et al. 2011).

What are the distinctive characteristics of this forest compared to other ecosystems where precipitation input has been experimentally altered? Contrary to our main hypothesis, we found no evidence of exclusion understory trees acclimate foliar traits to conserve o use more efficiently water. In fact, we found evidence of the opposite. There are two main reasons that may explain the behavior of individuals in the exclusion experiment. The first is the particular combination of the amount of rainfall and temperature in this temperate cool laurophyl forest. The southern South American temperate forests receive more than 2.000 mm. of rain every year. Our exclusion of throughfall removed a maximum of 13% of the rainfall (22% the last five years). This amount of water is still far from compromising the forest structure. (e.g. Whittaker 1975). Additionally, summer temperatures are not as high as those in the Mediterranean or tropical ecosystems (Park et al. 2019). Thus, trees do not show high transpiration levels during summer (i.e. a low VPD); consequently, water sources are not depleted at fast rates (Figure 7). For instance, flooded amazonian forests have traits related to a spender behavior, compared to non-flooded forests, possibly because the high VPD depletes soil water faster (Fontes et al. 2020). This is contrary to what we observed in this study. Second, the soils of Chiloé Island. Chiloé Archipelago has undergone a series of glaciations accompanied by volcanic activity (Moreno et al. 2021). These two processes constituted a soil that possesses a thin, but hard layer of aluminum oxides and iron deposits derived from volcanic activity (Díaz et al. 2007). This layer causes soil flooding, which drives plant species to handle occasional water-logging. Therefore, the decrease in soil moisture —observed at the start of measurements during the summer–autumn period of 2021 (see supplementary data)—associated with reduced soil water input in the exclusion plot, may have alleviated root hypoxia and/or anoxia in the treated individuals. The lower levels of ^δ^13C in the individuals used as controls may be evidence of this, as has been suggested in the bogged sites of *Metrosideros polymorpha* in Hawaii (Meinzer et al. 1992). Oxygen constraint alleviation resulted in increased performance of individuals who received less precipitation. Perez-Quezada et al. (2018), using eddy covariance data from this forest, may support this interpretation by showing that, at the ecosystem level, trees acquired more carbon during drier years compared to wetter ones.

**Figure 7.**
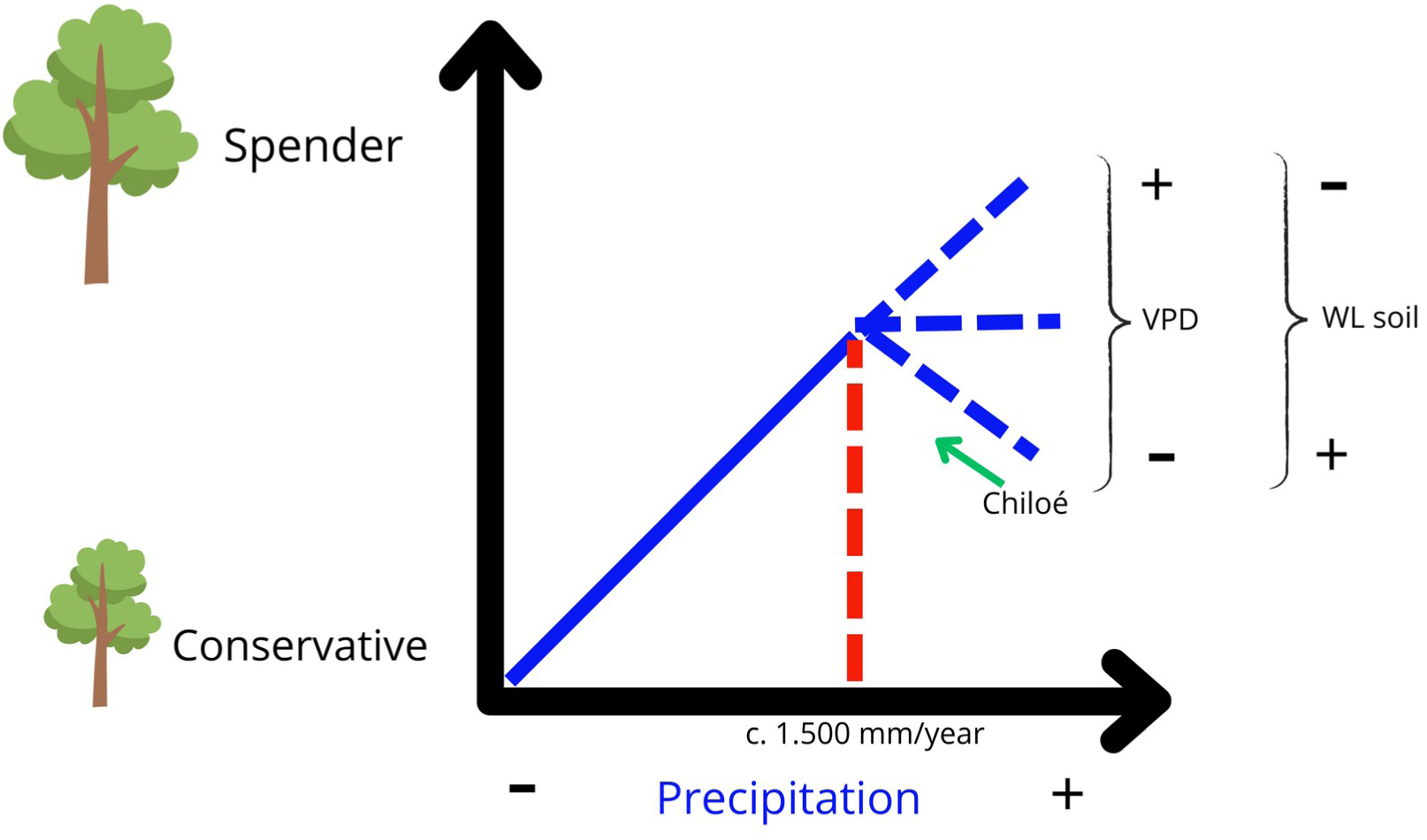
Acclimation model of trees due to precipitation, incorporating two key factors in Chiloé: Vapor pressure deficit (VPD) and waterlogged soils (WL). The red line represents the approximately level where trees just become more conservative (Derived from literature that address precipitation exclusion experiments in other forests). Average precipitation in Chiloé is c. 2.200 mm/year.

Considering that the species and individuals studied here are the future forest that would persist for centuries, these results have crucial implications for our predictive understanding of these ecosystems. If the tree species in the southern South America cold temperate forests, especially in Chiloé, are becoming more spender as a result of decreased rainfall input, this ecosystem may become in a large carbon sink the following centuries. Additionally, it would imply that spender trees may be less prepared to cope with droughts, when precipitation reductions stop to be an alleviation, as we registered here, and become a constrain (Fig. 7).

## Conclusions

To our knowledge, this is the first study on the manipulation of water input in a temperate cool laurophyl forest. Other studies assessing the effects of a precipitation experiment include climates with less precipitation (Mediterranean and temperate ecosystems) or higher temperatures (tropical forests). Evidence from this study shows that tree species have slightly acclimated at the leaf level because of a reduction in their water inputs. Nevertheless, this acclimation did not occur, as other similar studies have shown, and we first hypothesized. Trees, instead of being water conservatives, showed indications of becoming spenders. The differences were not equal for all species; however, most species exhibited changes in this direction. Even when the sum of the extracted water was low, some changes were observed. This suggests that higher reductions may acclimate the Chiloé forest to transpire more and consequently acquire more carbon. Although this possible increase in carbon gain with less precipitation is not sustainable, after a threshold of water reduction, this forest could become conservative or even not acclimate, as other forests have shown.

## Acknowledgments

We want to thank the Chilean National Agency of Research and Development (ANID), for the financing of this study, through the funds: Project FB210006, Fondecyt 1211765, and ANID Doctoral Scholarship 2018. We also want to thank Senda Darwin Biological Station for the support of this study. We respectfully acknowledge the invaluable contributions of Dr. Juan J. Armesto Z. to this study. He encouraged me to (Ben Castro) study the through-fall exclusion plots, actively participated in theoretical discussions and provided thoughtful revisions and guidance throughout the development of the work, up until his academic retirement. Regrettably, we were unable to publish this article prior to his passing.

## Author contributions

BC conceived the study, took the measurements, analyzed the data and wrote the manuscript. MFP was involved in theory discussion, analyzed data, revised the manuscript and was one of the researchers that installed the precipitation exclusion at Senda Darwin.

All authors have seen and approved the manuscript, and it hasn’t been accepted or published elsewhere.

## Data availability

All the data is available at reasonable request.

