## Supplementary information for "Physiological and anatomical leaf acclimation of understory trees subjected to a through-fall precipitation exclusion in a temperate rain forest in southern South America"

Supplementary data

Table 1. Leaf mass area (gr/m2, average ± standard deviation) of every species, all individuals, and families (with more than one species) from the exclusion and reference treatments. Symbols (*,**,***) display differences in t-test at 0.1, 0.05, and 0.01, respectively.

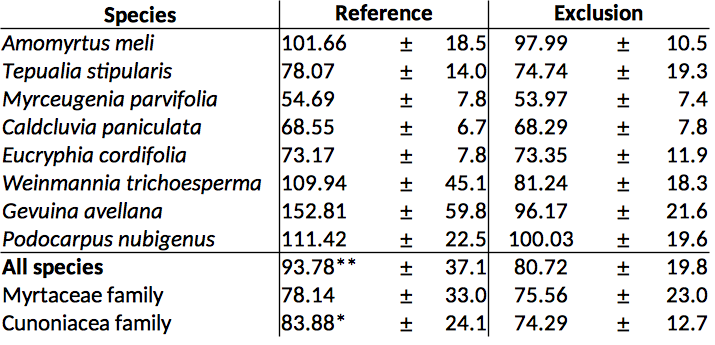

Table 2. δ13C, nitrogen, carbon, and C:N ratio for every species, all species, and families from exclusion and reference ecosystems. Symbols (*,**,***) displayed for differences in t-test at 0.1, 0.05 and 0.01 significance respectively.

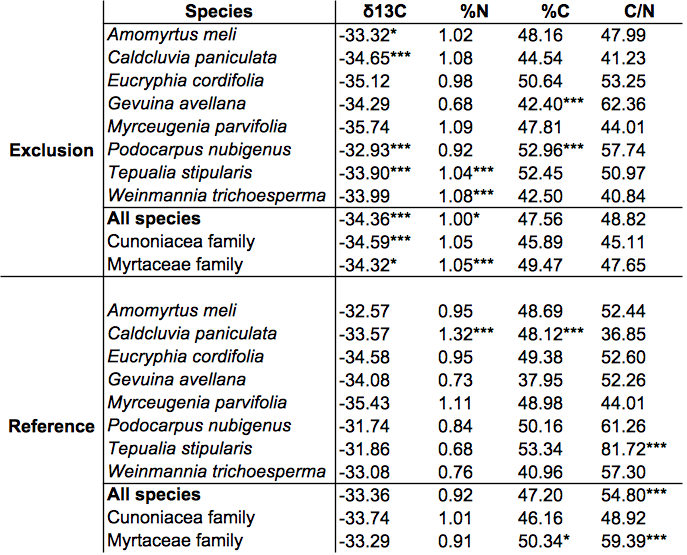

Table 3. Pressure-volume curve parameters of the eight species in the treatments. Average value ± standard error. Symbols (*,**,***) display differences in t-test at 0.1, 0.05, and 0.01, respectively.

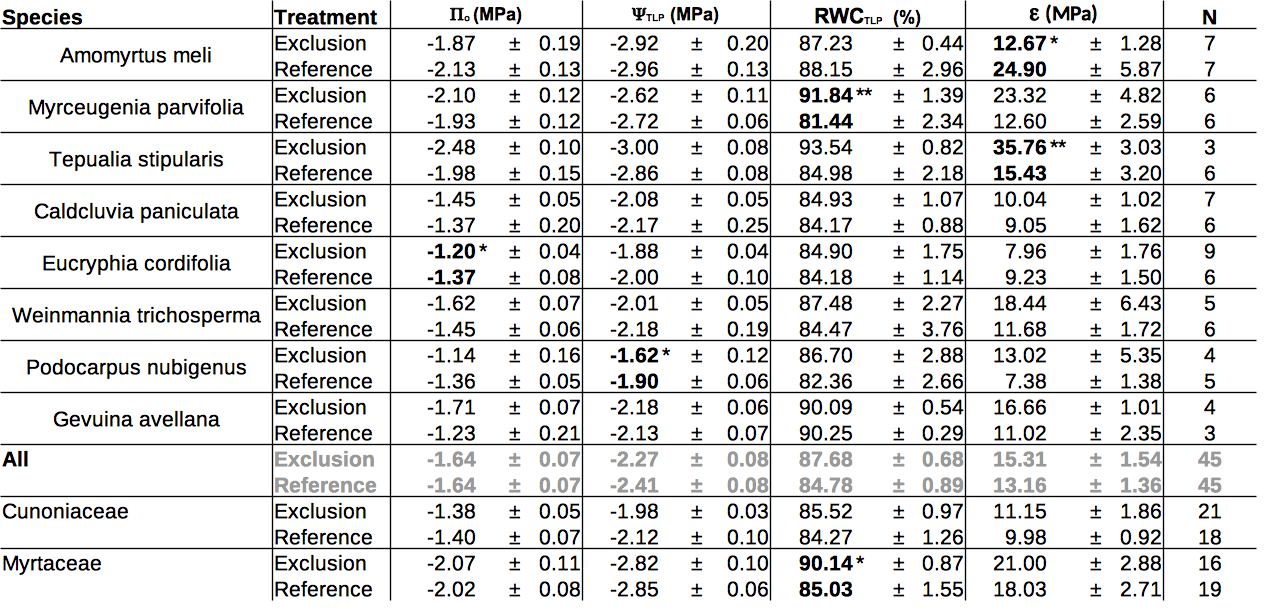

Table 4. Midday water potential by species for each treatment.

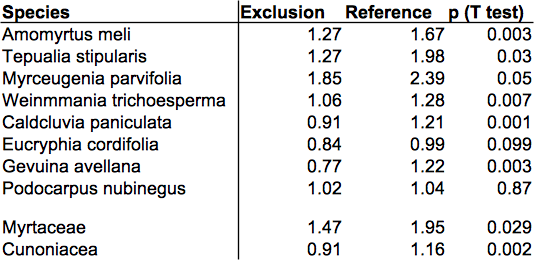

Graph 5. Gravimetric soil water content during summer – autumn 2021 (driest period) in both an exclusion plot and a reference soil.

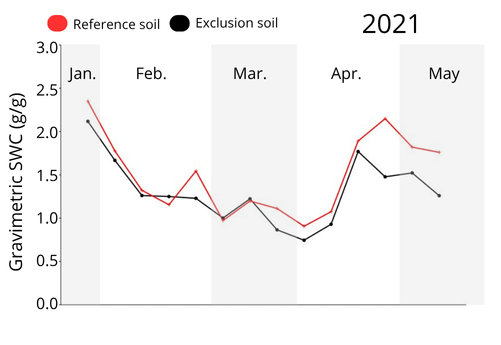

Table 6. Two way ANOVA (repeated measures, nine samples, 14 dates)

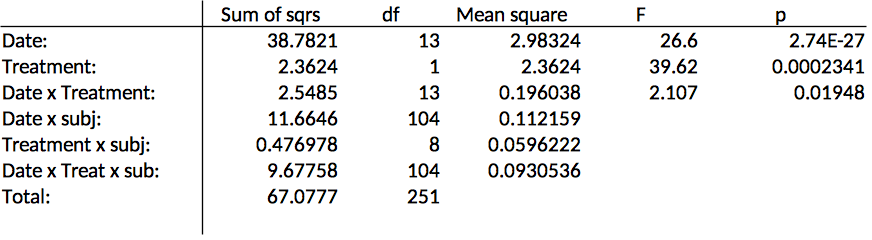
